# A Rare Nestin-Expressing Granule Cell Precursor Subpopulation Underlies SHH Medulloblastoma Formation

**DOI:** 10.64898/2026.05.26.727818

**Authors:** Salsabiel El Nagar, Yinwen Liang, Daniel N. Stephen, Synphen H. Wu, Alexandra L. Joyner

## Abstract

Granule cell precursors (GCPs) drive the major postnatal expansion of the cerebellum and are the cells of origin of sonic hedgehog medulloblastoma (SHH MB). Although GCPs are often treated as a uniform population, increasing evidence suggests they are heterogeneous, and whether specific subpopulations show distinct tumorigenic competence remains unclear. Here, we identified a rare Nestin-expressing GCP subpopulation in the normal early postnatal cerebellum and showed that it is spatially restricted, molecularly distinct and highly competent to form tumors. These cells are enriched in the posterior external granule layer and co-express *Atoh1*. Using different SHH MB mouse models, we showed that when this rare subpopulation of GCPs is targeted they can give rise to SHH MB with an efficiency comparable to targeting a larger number of *Atoh1*-expressing GCPs and that tumors derived from Nestin-expressing GCPs arise in the posterior-lateral cerebellum. Single cell RNA sequencing revealed that Nestin-expressing GCP have a transcriptome indicating reduced neuronal differentiation and enrichment for stem cell genes compared to bulk GCPs and more closely align with SHH MB cells. Together, our findings reveal functionally important heterogeneity within the GCP lineage and suggest that SHH MB arises preferentially from a small subpopulation of GCPs that express Nestin.

## Introduction

During cerebellar development, granule cell precursors (GCPs) drive much of the massive postnatal expansion and foliation of the cerebellum (Roussel and Hatten, 2011; Sudarov and Joyner, 2007). These progenitors, defined by the expression of *Atoh1*, originate from the rhombic lip germinal area, migrate to form a transient external granular layer (EGL) cell population on the surface of the cerebellum, and then undergo extensive expansion through cell division, before migrating inward and differentiating into mature granule neurons (Espinosa and Luo, 2008; Machold and Fishell, 2005; Wingate and Hatten, 1999). Traditionally considered a homogeneous population, GCPs have more recently been shown to exhibit molecular and functional heterogeneity (Consalez et al., 2021; Espinosa and Luo, 2008; Li et al., 2013; Wojcinski et al., 2019). However, the biological significance of the diversity remains poorly understood.

Nestin (NES), a type VI intermediate filament protein, is commonly used to label neural progenitors derived from the ventricular zone (VZ), including the VZ of the cerebellum, its second germinal area (Buffo and Rossi, 2013; Milosevic and Goldman, 2004). Indeed, in the postnatal cerebellum under normal conditions, *Nes* expression is restricted to neural and glial progenitors in the white matter and Bergmann glial layer referred to as NEPs (*Nes*-expressing progenitors) and not detected in the EGL (Wojcinski et al., 2017; Pakula et al., 2025). However, one set of studies (Li et al., 2013, 2016) identified a rare population of *Nes* positive (*Nes*^+^) cells under the EGL that exhibit GCP-like behavior and suggested that a subset of ventricular zone derived progenitors may transiently acquire granule progenitor features, even in the absence of pathological conditions. Another study also provided evident for *Atoh1*-expressing cells arise from the cerebellar VZ (Khouri-Farah et al., 2022). Thus, although most studies assume that *Nes* expression is restricted to non-rhombic lip lineages in the developing cerebellum, emerging evidence indicates that rare *Nes*⁺ cells might exist near the EGL, some of which potentially correspond to a small GCP subpopulation. Whether such cells exist during normal development and how they contribute to cerebellar lineages or cancer remains unclear.

This question is particularly important given that GCPs are the cell of origin for the sonic hedgehog (SHH) subtype of medulloblastoma (SHH MB), the most common pediatric malignant brain tumor) (Roussel and Hatten, 2011; Smith et al., 2022). In mouse models, GCPs are susceptible to MB formation when mutations result in the SHH pathway remaining constitutively active. Yet, it is unclear whether all GCPs share equivalent tumorigenic potential, or whether specific subpopulations exhibit unique vulnerabilities, perhaps distinguished by their spatial localization or molecular identity.

Here, we identified and characterized a rare subpopulation of *Nes-*expressing (*Nes^+^*) GCPs within the EGL during normal postnatal cerebellar development. These cells are specifically enriched in the posterior EGL at postnatal day 1 (P1), co-express the rhombic lip derivative marker *Atoh1* and exhibit a lower proliferation rate compared to *Nes* negative (*Nes*^-^) GCPs. Single cell RNA-sequencing (scRNA-seq) analysis provided additional evidence that *Nes^+^* GCPs are more quiescent and less differentiated than *Nes*^-^ GCPs and more closely related to GCP-like cells found in SHH MB (Tan et al., 2018; Lao et al., 2026). Using lineage tracing and SHH MB mouse models, we demonstrated that *Nes*⁺ GCPs are competent to give rise to SHH MB with a higher frequency than the general GCP population when targeted to express an activated form of the SHH receptor Smoothened (SMO). Furthermore, tumors arising from the *Nes*-lineage form preferentially in the posterior and lateral region of the cerebellum, corresponding to where SHH MB is commonly seen in mouse and human. Our findings therefore reveal a novel GCP subpopulation defined by their spatial and molecular identity, that could represent the main cell-of-origin of SHH MB.

## Results

### A rare subpopulation of *Nes*-expressing granule cell precursors is spatially restricted to the posterior cerebellum

To investigate whether *Nes* is expressed in GCPs within the EGL, we analyzed cerebella from *Nes-CFP* transgenic mice at P1 expressing CFP from promoter/enhancer sequences from the rat *Nes* gene (Encinas et al., 2006). Immunofluorescence staining for CFP, BARHL1 (a GCP marker), and DAPI revealed a rare population of CFP^+^ BARHL1^+^ double-positive cells located specifically within the EGL (Figure 1A-D). Higher magnification images indicated that the double-positive cells were largely absent in anterior regions of the cerebellum (Figure 1A’–D’) and highly enriched in the posterior portion of the EGL (Figure 1A’’–D’’), highlighting a spatial restriction. Quantification of serial sagittal sections across the medial-lateral axis confirmed this posterior enrichment (Figure 1E, F). Indeed, double-positive CFP^+^ BARHL1^+^ cells were almost exclusively detected in posterior lobules (folds of the cerebellum) (square bracket in Figure 1A-D). Based on systematic cell counts across half of the cerebellum (n=5 mice), there were an average of 18,189 ± 181 CFP^+^ GCPs per cerebellum at P1. These results identify a distinct posterior enriched *Nes*^+^ (CFP+) GCP subpopulation within the EGL (Figure 1E-F). We next examined whether these cells co-express SOX2, a marker of neural progenitors and co-expressed with *Nes* (CFP) in the ventricular layer, molecular layer and white matter (Wojcinski et al., 2017; Bayin et al., 2021). In the posterior EGL, a subset of CFP^+^ BARHL1^+^ cells were found to overlap with SOX2^+^ cells (Figure 1G-L). Quantification revealed that while a majority of CFP^+^ BARHL1^+^ cells were SOX2^−^ (69%), a notable proportion co-expressed SOX2 (20%) (Figure 1K-L), indicating molecular heterogeneity within the *Nes*^+^ GCP population. Interestingly, 11% of BARHL1^+^ cells where SOX2^+^ CFP^-^.

**Figure 1.**
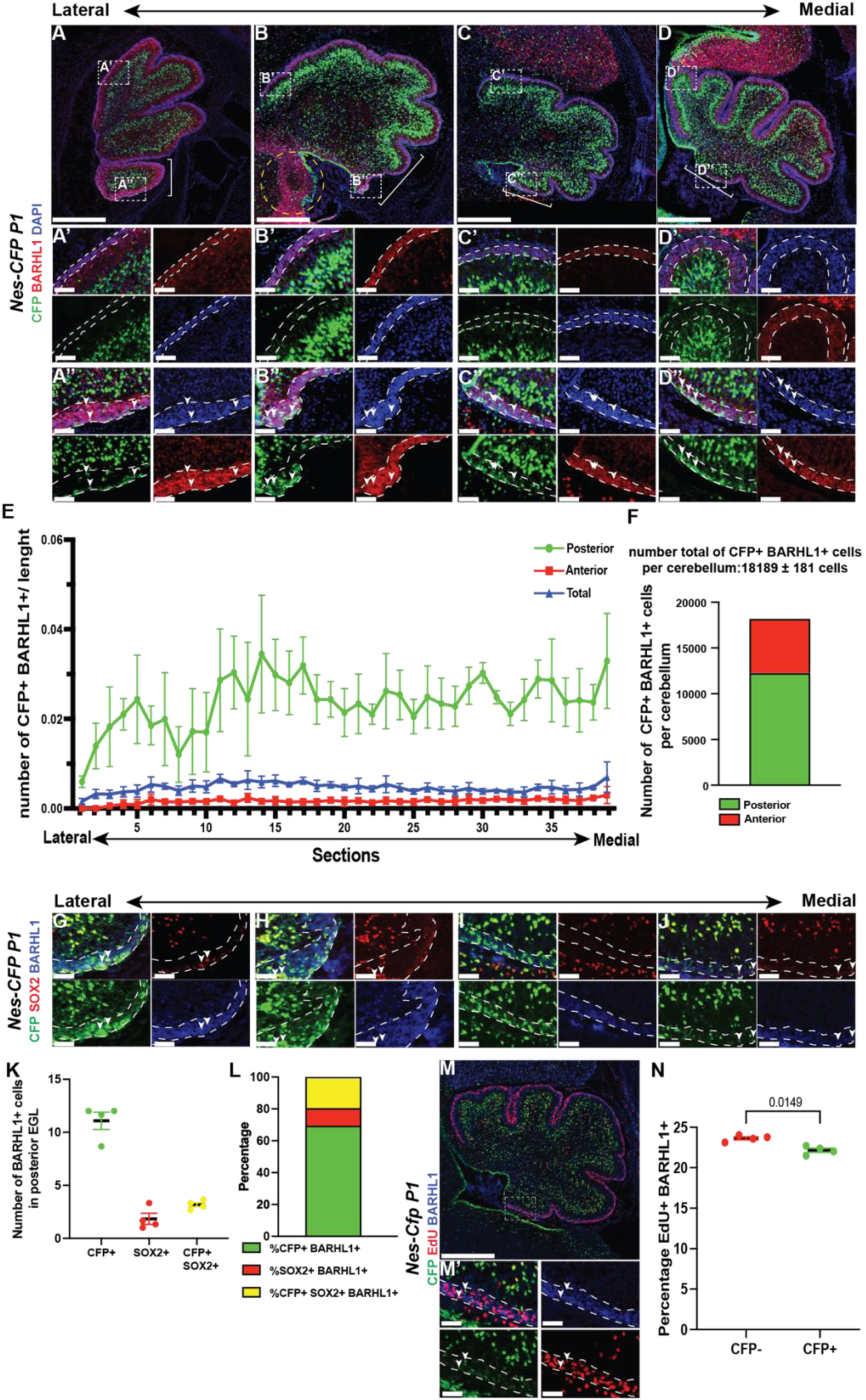
*Nes*-expressing granule cell precursors (GCPs) are enriched in the posterior cerebellum and display reduced proliferation. (A-D) Immunofluorescence staining of sagittal cerebellar sections from *Nes-CFP* mice at P1 showing CFP (green), BARHL1 (red) and DAPI (blue). Sections represent four distinct levels on the lateral-medial axis. Panels (A’-D’) and (A’’-D’’) show high-magnification images of anterior and posterior regions indicated by white dashed boxes in A-D. White arrowheads indicate examples of CFP^+^ BARHL1^+^ double positive cells. Yellow dashed circle indicates a cochlear nucleus. N>3 mice **(E)** Quantification of the CFP^+^ BARHL1^+^ cells normalized to EGL length along the lateral-medial axis in the anterior cerebellum and posterior cerebellum. The posterior cerebellum corresponds to the last cerebellar lobule (fold) delineated by white brackets in panel A-D, whereas the anterior corresponds to the remaining EGL. N=5 mouse samples. **(F)** Total number of CFP^+^ BARHL1^+^ cells per cerebellum at P1 in the posterior, anterior and entire cerebellar compartments. Total number of CFP^+^ BARHL1^+^ in the posterior is 12258 ± 106 cells and in the anterior is 5931 ± 76 cells. N=4 mouse samples. **(G-J)** Immunofluorescence staining of sagittal cerebellar sections from *Nes-CFP* mice at P1 showing CFP (green), SOX2 (red) and BARHL1 (blue). Panels show high-magnification images of the posterior cerebellum at four distinct lateral-medial levels. White arrowheads indicate some CFP^+^ SOX2^+^ BARHL1^+^ triple positive cells. N>3 mice. **(K)** Quantification of the number of CFP^+^ BARHL1^+^, SOX2^+^ BARHL1^+^ and CFP^+^ SOX2^+^ BARHL1^+^ cells per section in the posterior EGL of *Nes-CFP* cerebella at P1. N=4 mouse samples **(L)** Percentage of CFP^+^ BARHL1^+^ (69.2±4.2%), SOX2^+^ BARHL1^+^ (10.9±4.5%) and CFP^+^ SOX2^+^ BARHL1^+^ (19.8±1.3%) cells per in the posterior EGL of *Nes-CFP* cerebella at P1. N=4 mouse samples. **(M)** Immunofluorescence staining of a midsagittal cerebellar section from *Nes-CFP* mice at P1 showing CFP (green), EdU (red) and BARHL1 (blue). (M’) High magnification image of the posterior EGL. White arrowheads indicate some CFP^+^ EdU^+^ BARHL1^+^ the triple positive cells. **(N)** Quantification of the proportion of EdU^+^ cells within BARHL1^+^ CFP^-^ and BARHL1^+^ CFP^+^ populations. N=4 mousse samples. Statistical significance was determined using an unpaired t-test. Scale bar: A-D, M: 500µm; A’-D’, A’’-D’’, G-J, M: 50µm. Data are represented as mean ±SEM.

We next evaluated the proliferative status of the CFP^+^ BARHL1^+^ cells by administering a 1-hour pulse of EdU before euthanasia and stained sections for CFP, EdU and BARHL1 (Figure 1M). CFP^+^ EdU^+^ cells were observed in the posterior EGL (BARHL1^+^), confirming that some *Nes*-expressing GCPs are proliferative (Figure 1M’). Interestingly, quantification revealed that CFP^+^ BARHL1^+^ GCPs had a significantly lower percentage of EdU^+^ cells than their CFP^−^ BARHL1^+^ counterparts in the posterior lobules of the medial cerebellum (Figure 1N; p = 0.0149), suggesting a distinct proliferative behavior of *Nes*^+^ GCPs compared to the bulk of the population. Together, these data reveal a new, spatially restricted subpopulation of *Nes*-expressing GCPs in the posterior cerebellum that is distinct from the previously characterized CFP^+^ cells in *Nes-CFP* mice situated below the EGL since they express BARHL1 and they exhibit reduced proliferation compared to other GCPs.

### *Nes*-expressing GCPs co-express *Atoh1* and can be isolated by flow cytometry

We next asked whether *Nes*^+^ GCPs (CFP^+^) express the rhombic lip marker *Atoh1* by generating *Nes-CFP; Atoh1-GFP* (Rose et al., 2009) double transgenic mice, allowing us to label and isolate GCPs expressing the two fluorescent markers. Using flow cytometry analysis on cerebellar cells from *Atoh1-GFP*; *Nes-CFP* P1 mice, we identified four distinct cell populations: CFP⁻ GFP⁻, CFP⁺ GFP⁻, CFP⁻ GFP⁺, and CFP⁺ GFP⁺ (Figure 2A). Quantification of each cell population revealed that the double-positive population represents only a small fraction of total cerebellar cells at P1 (1.19%), whereas most cells were double-negative or single-positive for either GFP (*Atoh1^+^*) or CFP (*Nes*^+^ NEPs) (Figure 2B). This result confirms that the *Nes*^+^ GCP subpopulation is rare. To validate co-expression of *Nes* and *Atoh1* RNA, we sorted the four populations and performed RT-qPCR on each group. Expression of *Atoh1*, *Nes*, *Gfp*, and *Cfp* was measured in the single-or double-positive populations (GFP^+^ and/or CFP^+^) and compared to the double-negative cells (Figure 2C, Figure 2 – Source data 1). As expected, GFP^+^ cells showed strong *Atoh1* and *Gfp* expression but minimal *Nes*, whereas CFP^+^ cells expressed high levels of *Nes* and *Cfp*, but low *Atoh1*, indicating that the two single fluorescent populations were distinct. In contrast, the double-positive population co-expressed both *Atoh1* and *Nes* transcripts, along with their respective fluorescent reporters (Figure 2D, Figure 2 – Source data 1). Compared to double-negative cells, *Atoh1* and *Nes* were significantly higher in the double-positive fraction, supporting the identity of the cells as a previously unrecognized rare population of GCPs at P1 that expresses both the GCP marker *Atoh1* and ventricular zone marker *Nes*.

**Figure 2.**
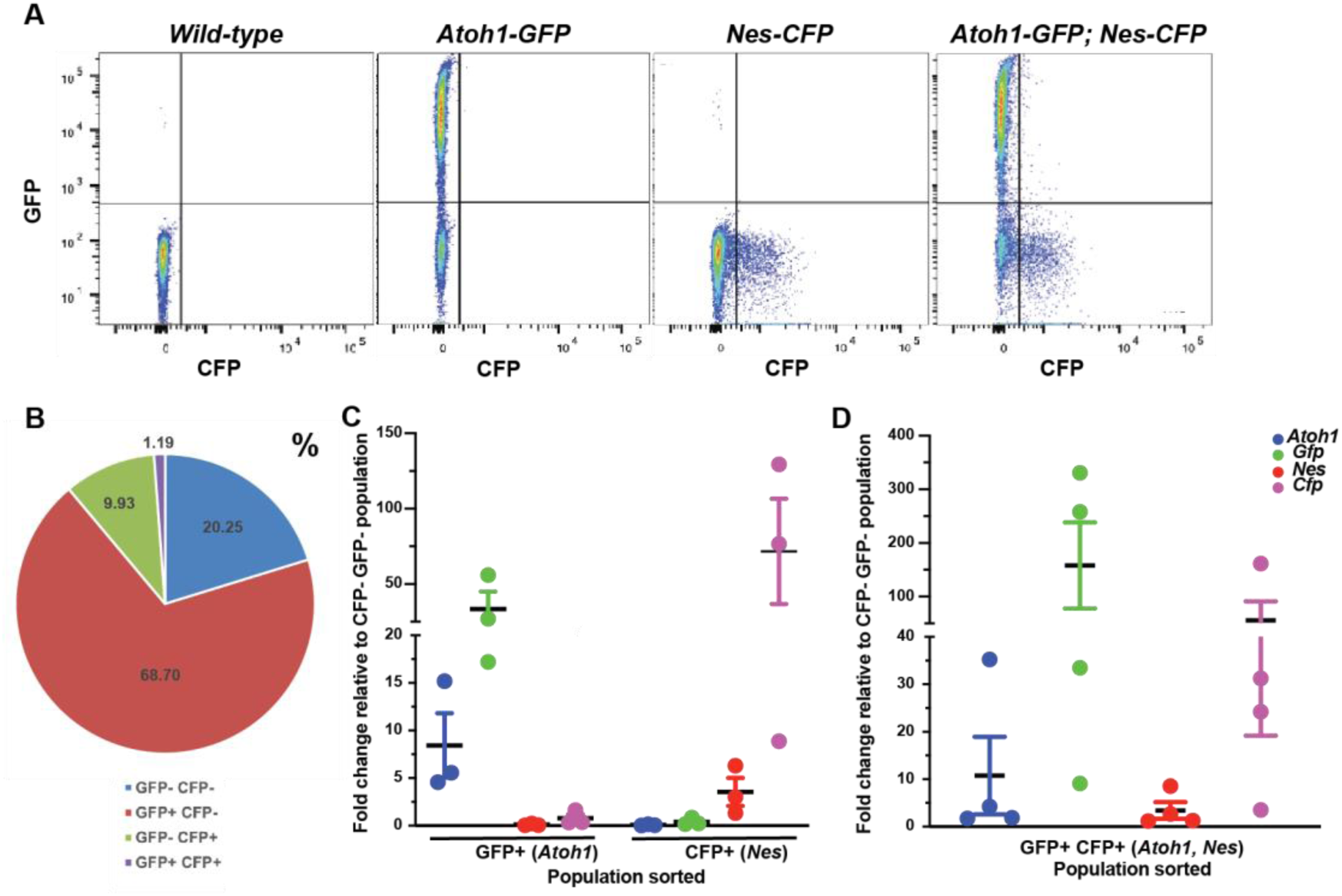
***Nes*-expressing GCPs can be isolated by flow cytometry and co-express *Atoh1*. (A)** Representative flow cytometry analysis of transgene negative (wild-type), *Atoh1-GFP* and *Nes-CFP* mouse cerebella at P1 used as controls and of *Nes-CFP; Atoh1-GFP* mouse cerebella to isolate the 4 cell populations GFP^-^ CFP^-^ (double negative), GFP^+^ CFP^-^ (*Atoh1*^+^ only), GFP- CFP+ (*Nes*^+^ only) and GFP^+^ CFP^+^ (double positive cells) in the *Atoh1-GFP; Nes-CFP* mouse cerebella. **(B)** Quantification of each population as a percentage of total live cells from P1 *Nes-CFP, Atoh1-GFP* cerebella. N=4 mouse samples. Percentages of each population ± SEM are: GFP^-^ CFP^-^: 20.25±7.87; GFP^+^ CFP^-^:68.70±8.55; GFP^-^ CFP^+^: 9.93±0.84; GFP^+^ CFP^+^: 1.19±0.23. **(C-D)** RT-qPCR analysis of sorted cell populations showing relative gene expression (fold change relative to GFP^-^ CFP^-^ cells). Panel (C) shows the single positive populations (GFP^+^ CFP^-^ and GFP^-^CFP^+^ cells) and (D) shows the double positive population GFP^+^ CFP^+^. N=3 and 4 biological replicates, normalized to *Gapdh*, respectively. See also Figure 2 – Source data 1. Data are represented as mean ±SEM.

### *Nes*-expressing GCPs are competent to form SHH MB

To determine whether *Nes*^+^ GCPs have tumorigenic potential, we tested their ability to form SHH MB. Two mouse models were generated that expresses Flp recombinase from either *Atoh1* or *Nes* promoter/enhancer sequences (Rose et al., 2009; Lao et al., 2012), along with two conditional alleles in the *Rosa26* (*R26*) locus referred to as SmoM2 mice (*R26^FSF-GFPcre/LSL-SmoM2^*). SmoM2 mice express Cre in Flp-expressing cells (*R26^FSF-GFPCre^* where FSF refers to frt-stop-frt) (Lao et al., 2012) and an activated form of SMO (SMOM2) in Cre expressing cells (*R26^LSL-SmoM2^* where LSL refers to lox-stop-lox) (Mao et al., 2006) (Figure 3A). Tamoxifen was injected at P0 to induce Flp activity which induced expression of a GFPcre fusion protein which then induces expression of SmoM2 and acts as a fluorescent reporter. A conditional loss-of-function allele of *Trp53* (Marino et al., 2000) was included in some mice to enhanced tumor penetrance (Figure 3B). Kaplan-Meier survival analysis revealed that *Atoh1-FlpoER; R26^/FSF-GFPcre/LSL-SmoM2^* (referred to as *Atoh1-*SmoM2) and *Nes-FlpoER; R26 ^FSF-Cre/LSL-SmoM2^* (referred to as *Nes-*SmoM2) mice both initiated tumorigenesis within 150 days upon constitutive SHH pathway activation, although *Atoh1-*SmoM2 mice showed a significantly shorter survival than *Nes-*SmoM2 mice (median survival 128.5 days vs. >150 days, p=0.0475)(Figure 3B). In the absence of *Trp53* in GCPs (*Atoh1-*SmoM*2; Trp53^fl/fl^* and *Nes-*SmoM2*; Trp53^fl/fl^*), both genotypes exhibited similar tumor incidence and shorter survival (medial survival 63.5 days for both) and showed that loss of *Trp53* strongly enhances tumor aggressiveness (Figure 3B). Histological analysis of tumors at P35, prior to overt morbidity, revealed distinct spatial patterns of lesion development based on genotype. In *Atoh1-*SmoM2*; P53^fl/fl^* mice, GFP^+^ lesions were distributed broadly across the surface of the cerebellum, consistent with the widespread presence of *Atoh1*^+^ GCPs throughout the EGL at P1 (Figure 3C-F). However, the largest lesions were detected along the surface of the posterior lobules mainly in the lateral cerebellum. In contrast, tumors arising in *Nes-*SmoM2*; P53^fl/fl^* mice exhibited more spatial restriction, with GFP^+^ lesions predominantly localized to the posterior and lateral cerebellum, corresponding to the more limited distribution of *Nes*^+^ GCPs (Figure G-J). These findings are similar to our previous finding that mouse SmoM2-stimulated SHH MB tumors induced with *Atoh1* transgenes only arises from the lateral cerebellum (Tan et al., 2018). At end-stage, medial and lateral sections from both models had large tumors that extended into the midline (Figure 3 K-N). Together, these results demonstrate that the rare *Nes*^+^ GCPs are competent to form SHH-driven MB and do so with temporal spatial dynamics that are distinct from bulk *Atoh1*^+^ GCPs.

**Figure 3.**
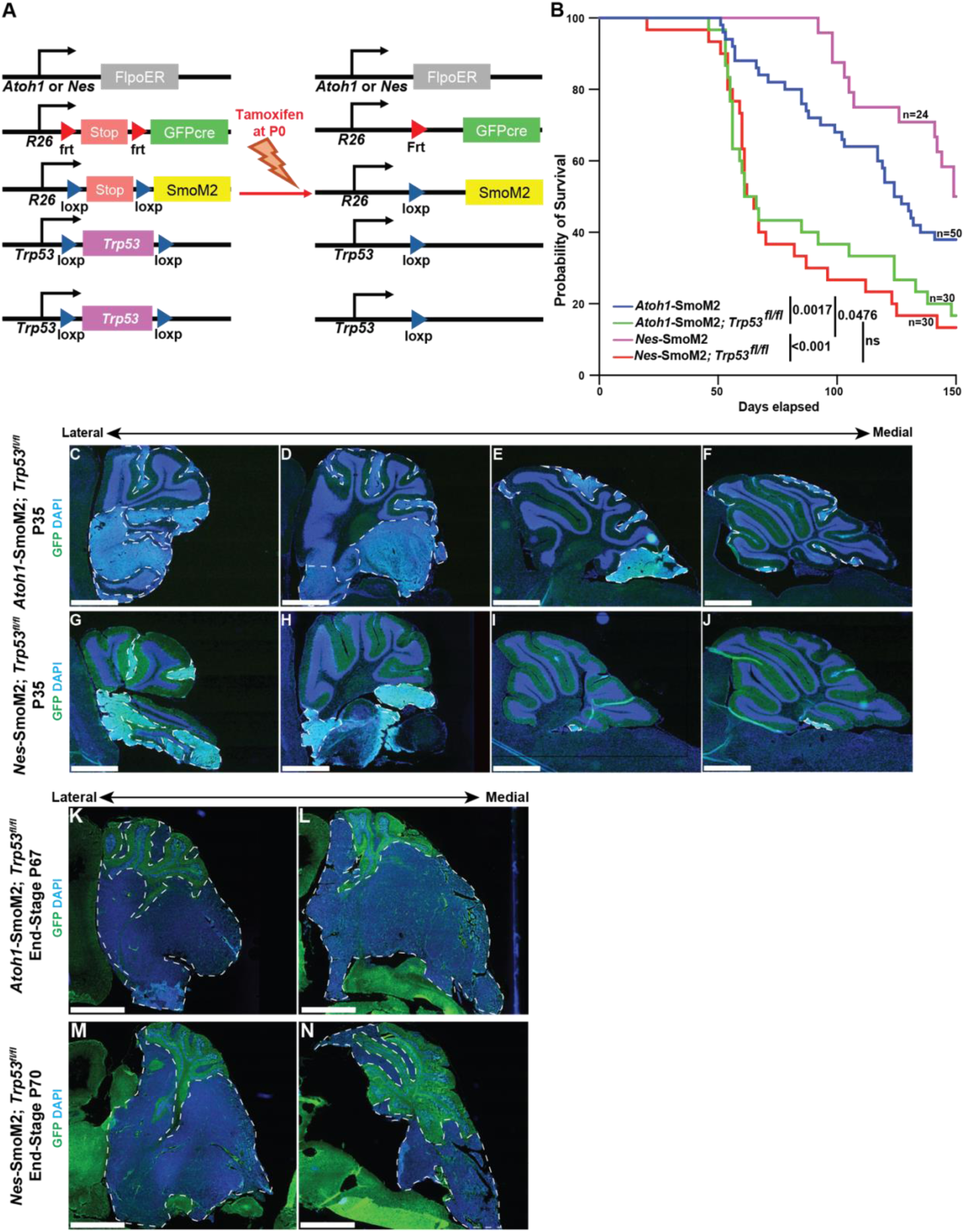
***Nes-*driven and *Atho1*-driven SHH medulloblastoma models show similar tumor penetrance and preferential growth of tumors laterally. (A)** Schematic representation of transgenes and gene loci in mosaic mutant tumor models *Atoh1-FlpoER; R26^FSF-GFPcre/SmoM2^; Trp53^fl/fl^* (*Atoh1-*SmoM2*; Trp53^flfl^*) and *Nes-FlpoER; R26^FSF-GFPcre/SmoM2^; Trp53^fl/fl^* (*Nes-*SmoM2*; Trp53^flfl^*) before and after tamoxifen injection at P0. **(B)** Kaplan-Meier curves of *Atoh1*- and *Nes*-driven mouse tumor cohorts with and without *Trp53* ablation: *Atoh1*-SmoM2, *Atoh1*-SmoM2; *Trp53^fl/fl^*, *Nes*-SmoM2 and *Nes*-SmoM2; *Trp53^fl/fl^*. Statistical significances were calculated using the log-rank test and P values are shown. **(C-J)** Immunofluorescence staining of sagittal cerebellar sections from an *Atoh1-*SmoM2*; Trp53^flfl^* (upper panel) and *Nes-*SmoM2*; Trp53^flfl^* (lower panel) mouse at P35 showing GFP (tumor cells, green) and DAPI (blue). Sections are showed at four distinct lateral-medial levels. Tumor lesions are delineated by white dashed lines. N>3 mouse samples. **(K-N)** Immunofluorescence staining of sagittal cerebellar section from an *Atoh1-*SmoM2*; Trp53^flfl^* (upper panel) and *Nes-*SmoM2*; Trp53^flfl^* (lower panel) mouse at end stage showing GFP (tumor cells, green) and DAPI (blue). Sections are showed in the hemispheres (lateral) and vermis (medial). Tumors are delineated by white dashed lines. N>3 mouse samples. Scale bar: C to N: 1mm.

### *Nes-FlpoER* labels a smaller population of GCPs than *Atoh1-FlpoER*

To compare the efficiency and specificity of recombination between *Atoh1*- and *Nes*-driven tumor models, we analyzed *Atoh1-FlpoER*; *R26^LSL-tdTom/FSF-GFPcre^*(*Atoh1-FlpoER; tdTom* reporter) and *Nes-FlpoER*; *R26^LSL-tdTom/FSF-GFPcre^* (*Nes-FlpoER;* tdTom reporter) (Daigle et al., 2018) mice at P2 following tamoxifen induction at P0. Immunofluorescence for RFP (tdTomato) revealed that the *Nes-FlpoER* line labeled fewer cells in the EGL than the *Atoh1-FlpoER* line, with recombination largely restricted to the posterior cerebellum (Figure 4A–H). Quantification of RFP⁺ cells across sagittal sections showed an average of approximately 2,920 ± 4102 cells per cerebellum in the *Nes-FlpoER* model compared to 25,480 ± 953 cells in the *Atoh1-FlpoER* model, representing an approximate ten-fold higher targeting efficiency using *Atoh1-FlpoER* (Figure 4I–J). These results confirm that the *Nes-FlpoER* driver targets a rare and spatially restricted subpopulation of GCPs within the developing cerebellum.

**Figure 4.**
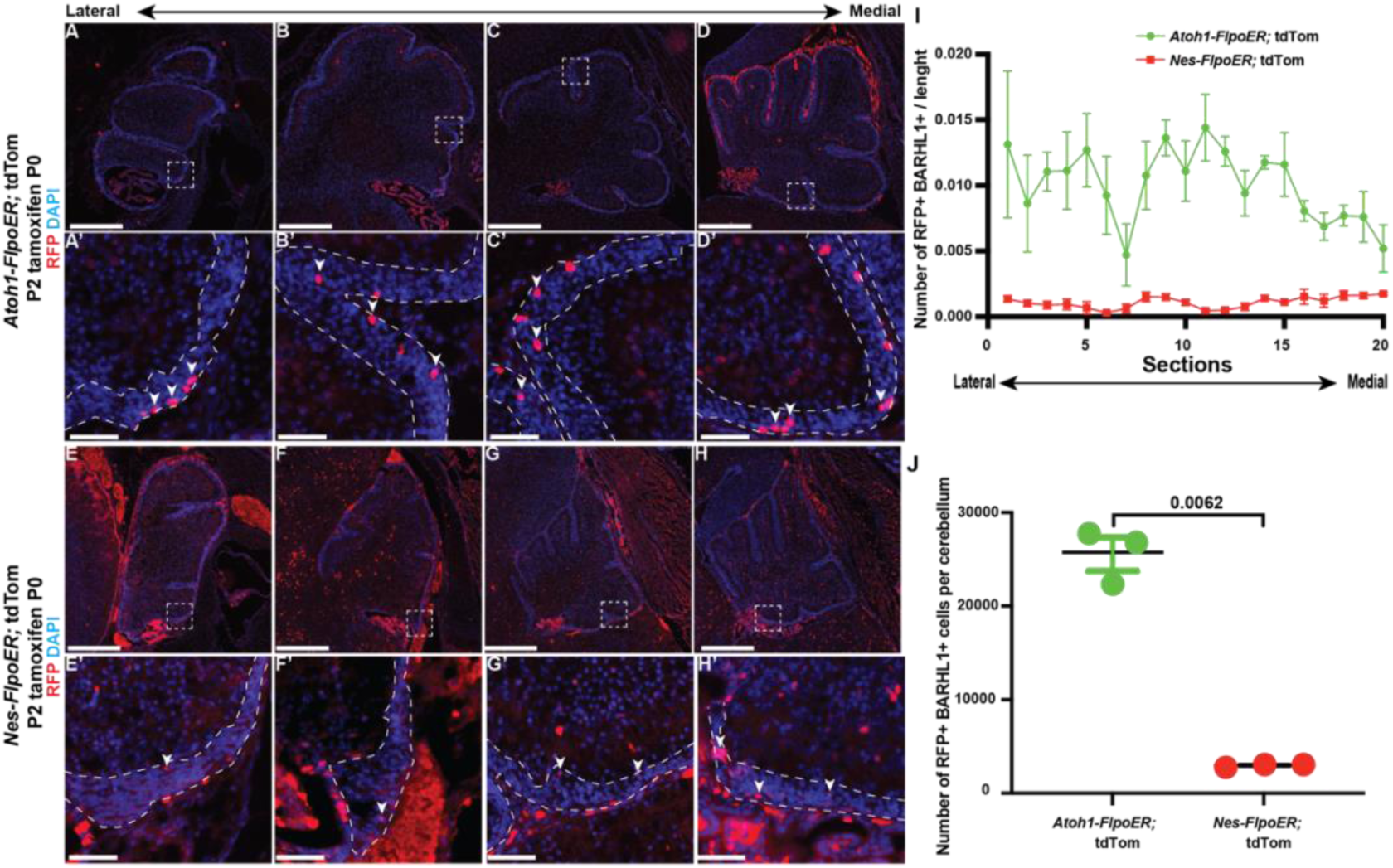
The *Nes*-driven SHH MB model targets approximately ten times fewer cells than the *Atoh1*-driven model. (A-D) Immunofluorescence staining of sagittal cerebellar sections from *Atoh1-FlpoER;* tdTom mice collected at P2 following tamoxifen injection at P0, showing RFP (red) and DAPI (blue). Sections are shown at four distinct lateral-medial levels. (A’-D’) High-magnification images of boxed regions in panels A-D. White arrowheads indicate the tdTomato^+^ cells (RFP) in the EGL. N>3 mouse samples. **(E-H)** Immunofluorescence staining of sagittal cerebellar sections from *Nes-FlpoER;* tdTom mice collected at P2 following tamoxifen injection at P0, showing RFP (red) and DAPI (blue). Sections are shown at four distinct lateral-medial levels. (E’-H’) High-magnification images of boxed regions in panels (A-D). White arrows indicate the tdTomato^+^ cells (RFP) in the EGL. N>3 mouse samples. **(I)** Quantification of the number of tdTomato^+^ (RPF) cells in the EGL along the lateral-medial axis in *Atoh1-FlpoER;* tdTom and *Nes-FlpoER;* tdTom mouse cerebella at P2 following tamoxifen injection at P0. N=3 mice. **(J)** Quantification of the number total of tdTomato^+^ cells in the EGL per cerebellum in both genotype *Atoh1-FlpoER; tdTom* and *Nes-FlpoER; tdTom.* N=3 mice. Statistical significance was determined using an unpaired t-test. Scale bar: A-D, E-H: 500um and A’-D’, E’-H’: 50um. Data are represented as mean ±SEM.

### *Nes-FlpoER* labels *Atoh1*-expressing cells capable of forming SHH MB

To validate that the targeted population corresponds to *Nes*⁺ GCPs and to reassess their ability to initiate MB, we developed an intersectional dual-recombinase model (*Nes-FlpoER; Atoh1^FSF-Cre/+^; R26^LSL-SmoM2/+^*) (Ruffault et al., 2015). In this model, oncogenic activation of SHH signaling occurs only in GCPs co-expressing *Atoh1* and *Nes*. Following tamoxifen induction at P0, these mice developed SHH MB with a latency and penetrance that seems comparable to those observed in the single-driver models (Figure 4–figure supplement 1A). Immunofluorescence analysis of *Atoh1^FSF-Cre/+^; Nes-FlpoER; R26^LSLtdTom/+^* cerebella collected at P4 confirmed RFP labeling within the EGL that displayed a pattern consistent with the spatial distribution of *Nes*⁺ GCPs (Figure 4–figure supplement 1B–E). Together, these findings validate that the *Nes-FlpoER* model effectively and selectively targets a minor subset of GCPs that co-express *Nes* and *Atoh1* and demonstrate that this restricted population is fully competent to initiate SHH-driven MB. In contrast, the *Nes-CreER* transgene used in many publications to model SHH MB does not match the endogenous *Nes* expression, since it targets many more GCPs in the EGL and throughout the anterior and posterior cerebellum compared to our *Nes-FlpoER* transgene, as shown by analyzing *Nes-CreER; R26^LSL-tdTom/+^* mice (Figure 4 – figure supplement 2).

### *Nes*-expressing GCPs exhibit a transcriptome associated with rhombic lip cells and quiescent stem cell identity

To characterize the molecular identity of the *Nes*⁺ GCP population we have identified, single-cell RNA-seq analysis was performed on five samples of P1 cerebella. Four independent samples were enriched for posterior GCPs using a Percoll gradient method and isolating the tissues from the posterior cerebellar region of individual *Nes-CFP* mice (Figure 5 – figure supplement 1A-C). A fifth sample contained CFP^+^ GFP^+^ cells isolated by FACS (fluorescence activated cell sorting) from the cerebella of 3 *Atoh1-GFP*; *Nes-CFP* mice. After filtering out low-quality cells (see Materials and Methods), the integration of 27,037 cells (22,961 posterior GCP-enriched samples and 4,076 CFP^+^ GFP^+^ cells) revealed 29 transcriptionally distinct clusters, including 12 GCP clusters expressing *Barhl1* (c0, c1, c2, c3, c7, c8, c9, c10, c13, c16 and c23) and 10 clusters with less cells expressing *Nes* (c4, c12, c14, c15, c17, c18, c19, c20, c21 and c27) that correspond to the ventricular zone-derivative NEPs (Figure 5A–C, Figure 5-source data 1; Figure 5-figure supplement 1D; Figure 5-figure supplement 1-source data 1). Additional clusters 11 and 22 represent unipolar brush cells, and five additional clusters likely represent degraded cells (5), fibroblasts (22), oligodendrocytes (24), microglia (25) and endothelial cells (26).

**Figure 5.**
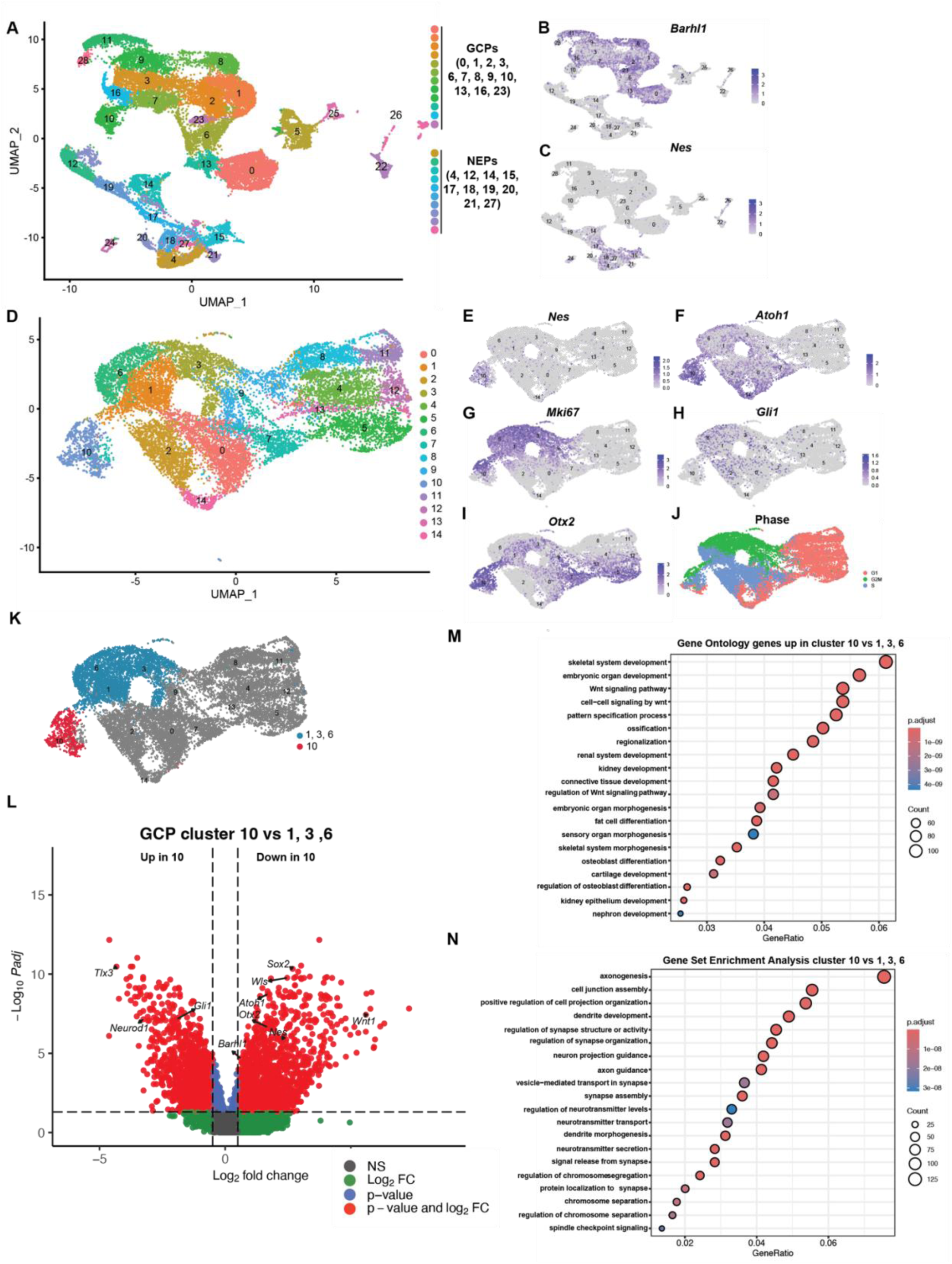
Single cell RNA-seq analysis identifies a *Nes*-expressing GCP subpopulation with distinct transcriptional features. **(A)** UMAP visualization of 5 samples of P1 cerebella cells showing 29 clusters. 4 samples were cells isolated from the posterior cerebellum and enriched from GCPs from individual mice and 1 sample (n=3 pooled) was GFP^+^ (*Atoh1^+^)* CFP^+^ (*Nes*^+^) cells isolated by FACS. *Atoh1*-expressing GCP clusters (0, 1, 2, 3, 6, 7, 8, 10, 13) and *Nes*-expressing progenitor (NEP) clusters (4, 12, 14, 17, 18, 19, 21, 22) are indicated. See also Figure 5-Source data 1. **(B–C)** Feature plots showing expression of *Barhl1* (GCP marker) and *Nes* across all clusters. **(D)** UMAP of integrated GCP clusters clusters (0, 1, 2, 3, 6, 7, 8, 10, 13 in part A) used for focused analysis. See also Figure 5 – Source data 2. **(E–I)** Expression of *Nes, Atoh1, Mki67, Gli1* and *Otx2* across the integrated GCP dataset. See also Figure 5-Source data 2. **(J)** Cell-cycle phase assignment showing G1, S, and G2/M populations in integrated GCP dataset. See also Figure 5-Source data 3. **(H)** UMAP highlighting in red cells in cluster 10 of the integrated GCP dataset that represent *Nes*+ GCPs and in blue cells in clusters 1, 3 and 6 expressing posterior cerebellum GCP markers. Grey indicates remainder of cells. **(L)** Volcano plot showing differentially expressed genes between cluster 10 and clusters 1, 3, and 6 in the integrated GCP dataset. Genes enriched in cluster 10 (right) include *Sox2, Wnt1*, and *Wls* genes enriched in clusters 1, 3, and 6 (left) include *Gli1* and *Neurod1.* See also Figure 5-Source data 4, 5. **(M-N)** GO enrichment plots showing biological processes upregulated in cluster 10 compared with other GCP clusters (1, 3, 6) in the integrated GCP dataset, including pathways related to cell-cycle progression, DNA replication, and oxidative metabolism. See also Figure 5 – Source data 6, 7.

The clusters containing GCPs were re-clustered producing 15 clusters, of which one transcriptionally distinct population, cluster 10, co-expressed *Nes* and *Atoh1* and the mouse line reporters *Cfp* and *Gfp* (Figure 5D-F, Figure supplement 1E; Figure 5-Source data 2). This cluster also expressed the proliferation marker *Mki67*, the SHH target gene *Gli1* and posterior GCP marker *Otx2* (Figure 5F-I; Figure supplement 1E; Figure 5-Source data 2). Cell cycle phase analysis showed that cells at all phases were present in cluster 10 (G1, G2M, S) (Figure 5J, Figure 5-Source data 3). Three neighboring clusters (1, 3, and 6) appeared to display a close transcriptional similarity to cluster 10, including the expression of *Otx2* and *Tlx3* which confirms these GCPs are localized to the posterior cerebellum (lobules 8-10) that contains the anatomical domain of *Nes⁺* GCPs (Figure 5I and Figure 5 – figure supplement 1E,F; Figure 5-Source data 2) (Divya et al., 2016; El Nagar et al., 2018).

Based on the similarity of the clusters, we compared the transcriptional profile of cluster 10 to the three posterior GCP clusters (Figure 5K). Comparison of cluster 10 (741 cells) with GCP clusters 1, 3, and 6 (4,066 cells) using the FindMarkers approach identified 1,586 genes upregulated in cluster 10 and 3,130 downregulated (Figure 5 - Source data 4) and using the more stringent Pseudobulk (Libra) analysis 1,983 upregulated and 1,876 downregulated genes (Figure 5L, Figure 5-Source data 5). Genes enriched in cluster 10 included *Sox2*, *Nes*, *Wls*, and *Wnt1*, while *Neurod1*, and *Rbfox3* were preferentially expressed in the other GCP clusters (1, 3, 6), indicating that cluster 10 represents a less differentiated, more stem-like GCP state, potentially positioned upstream in the lineage hierarchy.

Gene Ontology revealed that cluster 10 is enriched for biological processes linked to WNT signaling whereas genes associated with neuronal differentiation, synapse formation, and morphogenesis were downregulated (Figure 5M,N, Figure 5-Source data 6, 7). This enrichment of WNT-associated pathways and up regulation of *Wls* and *Wnt1* (Figure 5L), together with the fact that differentiation pathways are down regulated, suggests a transcriptional resemblance to rhombic lip progenitors. This molecular profile supports their identity as a transient, developmentally plastic population within the GCP lineage related to the rhombic lip.

### *Nes*-expressing GCPs are transcriptionally closer to GCP-like cells in SHH MB tumors than other GCPs

To further assess the relationship between the *Nes*-expressing normal GCPs and the GCP-like cells in SHH MB tumors, we integrated the cells of clusters 10, 1, 3 and 6 from our GCP cells (Figure 5K) with the GCP-like tumor cells from SmoM2 early tumors in a dataset we reported in Lao et al., 2026 (Lao et al., 2026) (Figure 6A-B, Figure 6- - data 1). We then performed pseudobulk differential expression analyses comparing GCP-like tumor cells with three GCP populations: all GCPs in the four clusters, *Nes*^-^ GCPs and *Nes*^+^ GCPs (Figure 6C-H).

**Figure 6.**
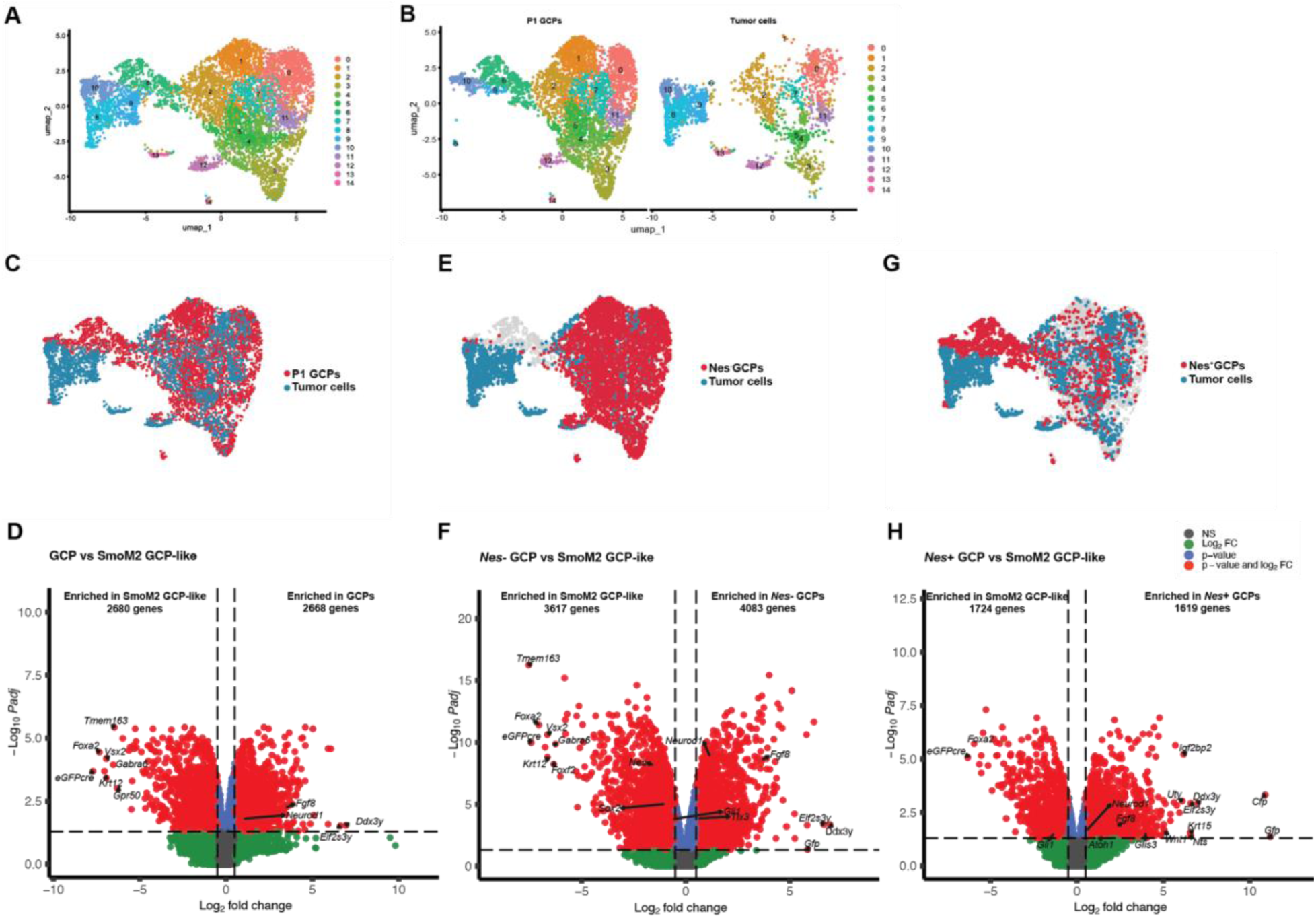
***Nes*-expressing GCPs appear transcriptionally closer to SHH MB tumor cells than *Nes*- GCPs. (A)** UMAP visualization of an integrated dataset of the GCP clusters 10, 1, 3 and 6 at P1 (see Figure 5D) with the GCP-like tumor cells from a SmoM2 model dataset (Lao et al., 2026). See also Figure 6-Source data 1. **(B)** UMAP showing the same integrated dataset as in (A) colored by sample of origin, distinguishing P1 GCP cells and SmoM2 GCP-like tumor cells. **(C, E, G)** UMAP highlighting in blue the SmoM2 GCP-like tumor cells and in red all GCPs (C), *Nes*^-^ GCPs (E) or *Nes*^+^ GCPs (G). **(D, F, H)** Volcano plots showing differential gene expression of all GCPs (D), *Nes*^-^ GCPs (F) or *Nes*^+^ GCPs (H) compared to SmoM2 GCP-like tumor cells. See also Figure 6-Source data 2-4.

Comparison of all GCPs with SmoM2 GCP-like tumor cells identified 2,680 and 2,668 significantly differentially downregulated and upregulated genes, respectively (Figure 6C-D, Figure 6-Source data 2). Comparison between *Nes*- GCPs and tumor cells identified 3,617 significantly downregulated genes and 4,083 significantly upregulated genes (Figure 6E-F, Figure 6-Source data 3). In contract, a comparison between *Nes*^+^ GCPs and tumor GCP-like cells yielded substantially fewer differentially expressed genes, with 1,724 and 1,619 significant downregulated and upregulated genes, respectively (Figure 6G-H, Figure 6 – Source data 4). Thus, among the three GCP population analyzed, *Nes*-expressing GCPs displayed the smallest transcriptional difference from the SmoM2 tumor cells. These data indicate that the *Nes*-expressing GCP population in the normal cerebellum is transcriptionally closer to SHH MB tumor cells than the remainder of the GCPs (clusters 1, 3, 6).

## Discussion

In this study, we identify a rare subpopulation of *Nes*-expressing GCPs in the normal developing cerebellum and showed that these cells are spatially restricted, molecularly distinct from other GCPs, and highly competent to form SHH MB. Although GCPs have long been considered the cells of origin of SHH MB, our findings refine this view by demonstrating that not all GCPs are equivalent in their propensity to form tumors. Instead, a minor *Nes*-expressing subset of GCPs within the EGL displays unique biological properties, including preferential posterior location, reduced proliferation, enhanced progenitor-like transcriptional characteristics and a high potential to form SHH MB. Together, these results reveal a new aspect of the heterogeneity within the GCP population and suggest that this heterogeneity is functionally relevant to both normal and tumoral cerebellar development.

An important feature of the *Nes*-expressing GCP population is its restricted localization to the posterior cerebellum. Previous work showed that mouse SHH MB preferentially arises from the posterior and lateral mouse cerebellum under conditional activation of SHH signaling and also human SHH MBs shows a similar anatomical preference (Teo et al., 2013; Wefers et al., 2014; Perreault et al., 2014; Zhao et al., 2017; Tan et al., 2018). Why only the laterally located *Nes*^+^ GCP give rise to tumors remains to be determined. Our data nevertheless support the idea that the regional enrichment of *Nes*-expressing GCPs may contribute to the known spatial preference of SHH MB formation. It is also interesting that CFP^+^ (*Nes^+^*) BARHL1^+^ cells are detected in the cochlear region (Figure 1B), a region where more than 30% of SHH MB arise from in human (Grammel et al., 2012).

Our analysis further showed that the majority of *Nes*-expressing GCPs do not express the neural stem cell marker SOX2, with only a minority of them showing co-expression of CFP (*Nes*), BARHL1 and SOX2. This is an important distinction as SOX2^+^ progenitors were proposed as candidate cells of the origin in SHH MB (Selvadurai et al., 2020). Our findings suggest that the *Nes*-expressing GCP population identified here is not equivalent to the previously described SOX2^+^ progenitor population and may instead represent an additional and distinct cell of origin of SHH MB. It will therefore be important to directly compare the tumorigenic potential of *Nes*^+^ SOX2^+^ and *Nes*^+^ SOX2^-^ GCPs. Furthermore, the presence of a smaller SOX2^+^ fraction within the *Nes*-expressing GCPs suggests an internal heterogeneity consistent with the possibility that these cells have distinct developmental or functional states with respect to tumor formation.

One of the most significant findings of this study is that a GCP population targeted at a ten-fold lower frequency than the broader *Atoh1*^+^ GCP pool nevertheless gives rise to SHH MB with a similar penetrance. This observation argues that tumor susceptibility is not determined simply by the number of targeted GCPs, but rather by the intrinsic biological properties of specific GCP subtypes. This result provides direct functional evidence that developmental heterogeneity within the GCP population translates into differential tumorigenic potential.

Our findings also help clarify prior work based on *Nes*-expression driven tumor models. Earlier studies using a *Nes*-*CreER* transgene concluded that the *Nes*-lineage cells can give rise to SHH MB (Li et al., 2013), but the precise identity of the targeted population remained difficult to resolve because the *Nes-CreER* transgene used is broadly expressed throughout the EGL and in multiple progenitor populations during development (Figure 4 – figure supplement 2). In contrast, the *Nes-FlpoER* transgene strategy used here labels much fewer cells and a spatially restricted subset of cells within the EGL. Furthermore, our intersectional genetic experiments provide additional support that the tumor-competent population corresponds specifically to cells that co-express *Nes* and *Atoh1*. Thus, our model likely provides a more precise readout and definition of the GCP subset of interest within the broader *Nes*-lineage.

The *Nes* gene might represent more than a simple marker of a tumor-prone GCP population and could also contribute directly to its tumorigenic behavior. Although *Nes* is commonly used as a marker of immature neural states, previous work in SHH MB showed that NES can functionally promote tumorigenesis by maintaining SHH signaling (Li et al., 2016). *Nes*-expression was shown to increase during tumor formation, and *Nes*-depletion found to reduce proliferation, promote differentiation and suppress tumor growth through regulation of GLI3 processing. These findings raise the possibility that *Nes* is not simply labeling a permissive progenitor population but may itself support the oncogenic SHH pathway activity. In this context, our data suggest an important nuance: that although initially *Nes*-expressing GCPs in the medial and lateral cerebellum form lesions, ultimately only laterally located *Nes*-expressing GCPs form tumors. One interesting question is whether this finding is related to the level of NES protein present in the cells, perhaps being higher in lateral cells. *Nes*-expressing GCPs therefore might not possess a greater ability to initiate tumor formation, but rather a stronger capacity to remain in a tumorigenic state once transformation has been engaged. It will be important to address this idea directly in future studies using genetic loss-of-function approaches within the *Nes*-expressing GCP population.

Our scRNA-seq characterization of *Ne*s^+^ GCPs further supports their identity as a progenitor or stem-like cells. *Nes*-expressing GCPs exhibit lower EdU incorporation in vivo and, and in the scRNA-seq data they segregate as a distinct GCP cluster with features suggestive of reduced differentiation and increased developmental plasticity. Compared with neighboring posterior GCP clusters, *Nes*-expressing GCP cluster cells retains GCP markers while showing enrichment for progenitor-associated genes and pathways, including components linked to WNT pathways (including *Wls*) (Yeung and Goldowitz, 2017) and a relative reduction of neuronal differentiation. This combination of reduced proliferative activity and reduced differentiation suggests that these cells may exhibit quiescent and stem-like features reminiscent of cancer stem cells (Chen et al., 2016). Indeed, these features may enhance their competence to maintain the oncogenic state.

An additional observation supporting the relevance of this population is its relationship to tumor cells at the transcriptomic level. Relative to other GCP clusters, *Nes*-expressing GCPs show fewer transcriptional difference from SHH MB GCP-like cells, suggesting that they may lie closer to the tumor cells than do other GCPs. This result implies that their developmental program may require fewer transcriptional changes to transit into a tumorigenic state under constitutive SHH activation. These observations fit well with our developmental findings, as a progenitor population that is relatively undifferentiated and still relatively plastic might be expected to transform more easily that other GCPs.

In summary, our study identifies a rare *Nes*-expressing GCP population as a distinct population with the cells located in the lateral developing cerebellum being highly tumor competent. These findings refine the view of GCPs as the cell of origin of SHH MB by showing that tumor susceptibility is linked to the heterogeneity within the GCP population. Finally, our findings support a model in which SHH MB arises preferentially from rare *Nes*-expressing GCPs defined by their spatial, molecular and functional properties.

## Materials and Methods

### Key Resource Table

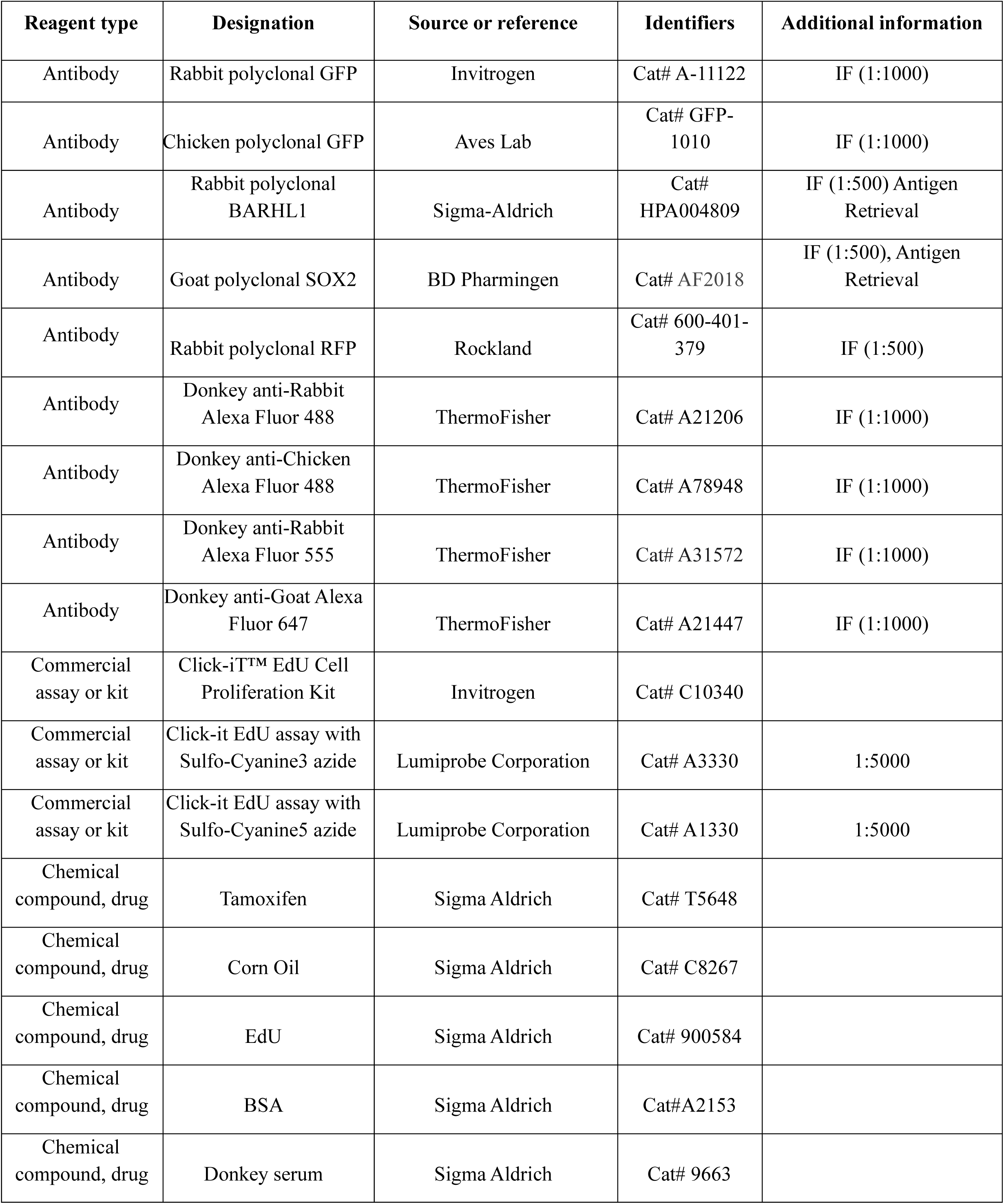
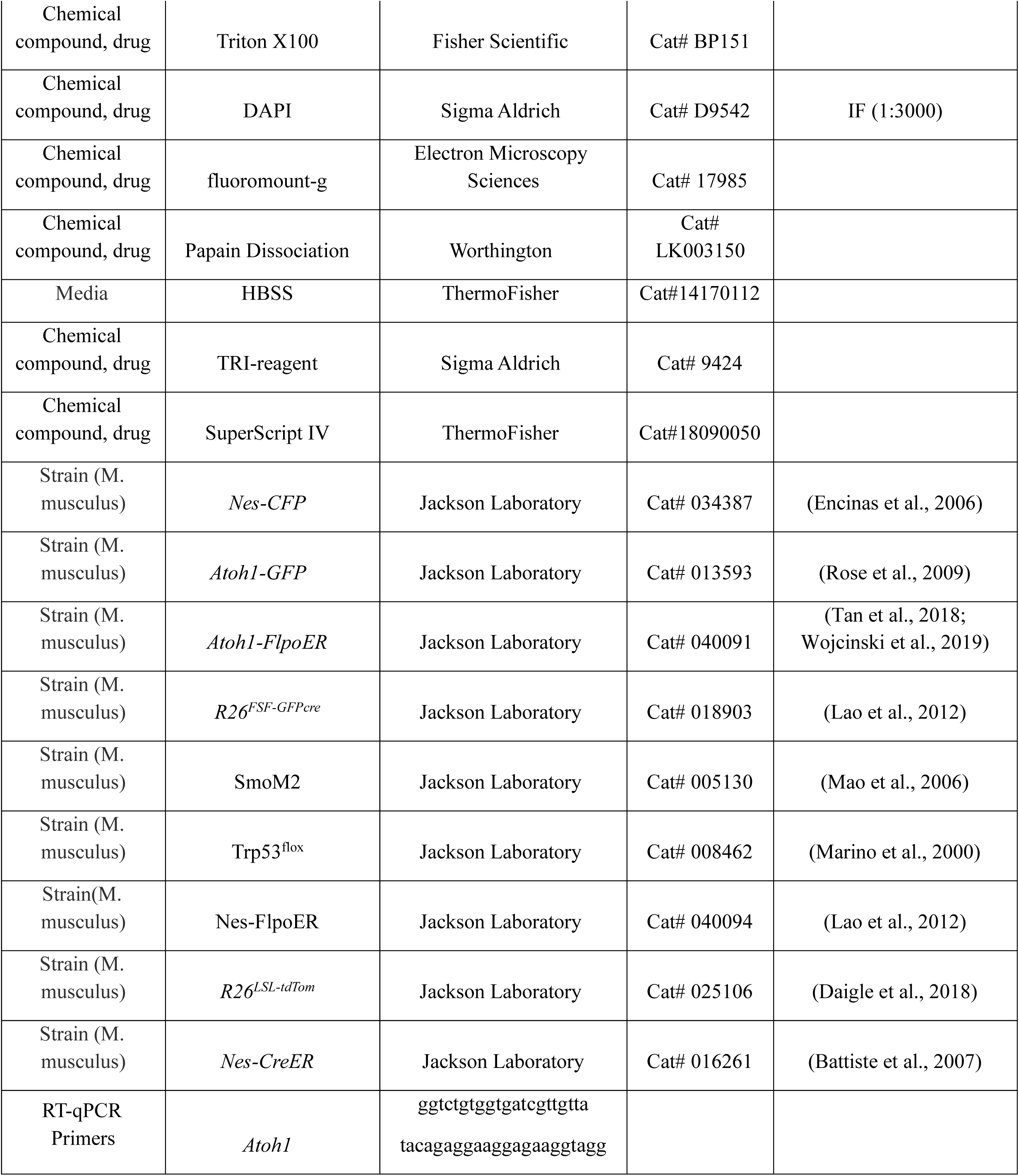
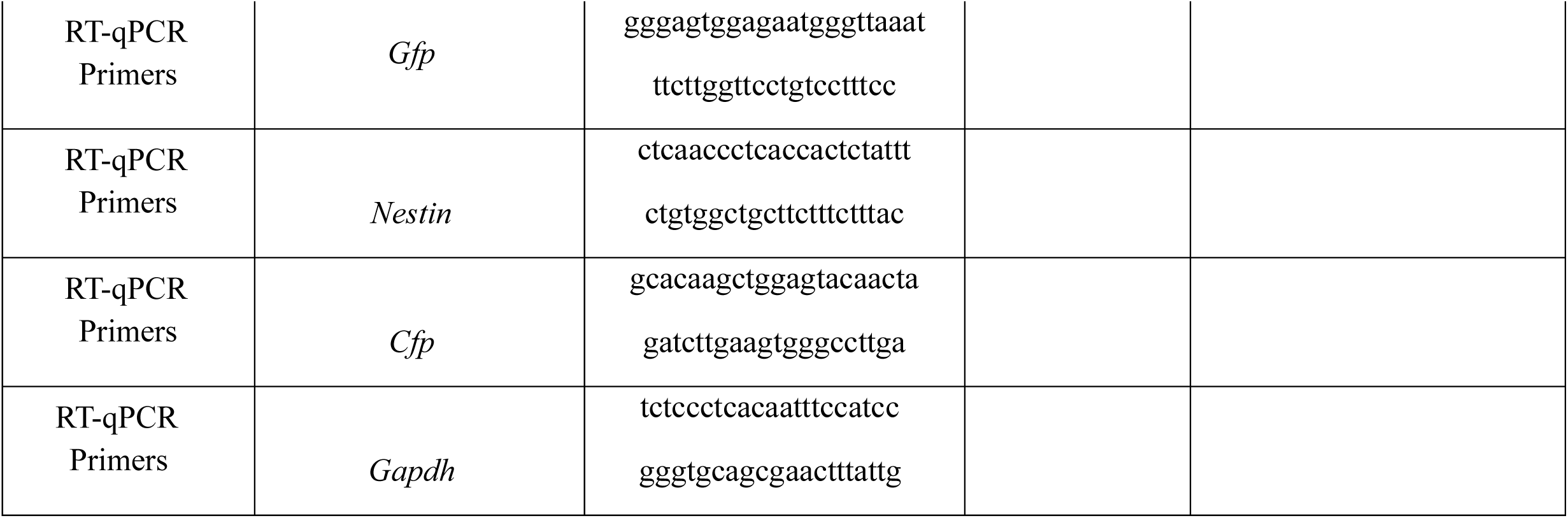

### Animal model and Breeding

All the mouse experiments were performed according to protocols approved by the Institutional Animal Care and Use Committee of Memorial Sloan Kettering Cancer Center (MSKCC) (protocol no. 07-01-001). Animals were housed in temperature-and humidity-controlled rooms on a 12-hour light/dark cycle and given access to standard laboratory mouse chow and water ad libitum. Both sexes were used randomly in this study. Experimenters were blinded for genotypes whenever possible. The following mouse line were used: *Nes-CFP* (JAX #034387) (Encinas et al., 2006), *Atoh1-GFP* (JAX#013593) (Rose et al., 2009), *Atoh1-FlpoER* (JAX #040091) (Tan et al., 2018; Wojcinski et al., 2019), SmoM2 (JAX #005130) (Mao et al., 2006), *R26^FSF-GFPcre^* (MASTR, JAX #018903) (Lao et al., 2012), *Trp53^flox^ (*JAX # 008462) (Marino et al., 2000), *Nes-FlpoER* (JAX #040094) (Lao et al., 2012), *R26^LSL-tdTom^* ( Ai75, JAX #025106) (Daigle et al., 2018), *Nes-CreER* (JAX #016261)(Battiste et al., 2007) and *Atoh1^FRT-Cre^* was obtained from Christo Goridis (Ruffault et al., 2015).

Tamoxifen administration: Genetic recombination in mouse lines was induced by injection of tamoxifen. Tamoxifen (T5648, Sigma-Aldrich) was dissolved in corn oil (C8267, Sigma-Aldrich) at 20mg/ml and injected subcutaneously into the back of P0 mice at a dose of 100mg per g of body weight.

EdU administration: 5-ethynyl-2’-deoxyuridine (EdU) (900584, Sigma-Aldrich) stock was dissolved in sterile phosphate-buffered saline (PBS) at 10 mg/mL and a dose of 5 μg/g was injected intraperitoneally into animals 1 hour prior to euthanasia.

### Tissue processing and immunostaining

For immunocytochemistry, animals younger than P8 were decapitated and then the whole head was fixed in 4% paraformaldehyde for 48 hours (hr) at 4°C, cryoprotected in 30% sucrose in phosphate-buffered saline (PBS) until they sank and then frozen in Cryo-OCT (Tissue-Tek). Older animals were anesthetized and then perfused with cold PBS followed by 4% paraformaldehyde prior to brain dissection. Frozen brains were sectioned sagittally at 12 μm or 14 μm and slides stored at −80°C.

For immunofluorescence staining, slides were left to warm to room temperature (RT) for 1 hr and washed 3 times in PBS. Primary antibodies diluted in the blocking buffer were placed on slides for overnight incubation at RT. The blocking solution was composed of 4% Bovine serum albumin (BSA) (A2153, Sigma Aldrich), 3% donkey serum ((D9663, Sigma Aldrich) and 2% Triton X100 (BP151, Fisher Scientific) dilatated in PBS. Slides were then washed 3 times in PBS and incubated with fluorophore-conjugated secondary antibodies diluted in the blocking buffer for 2 hr at RT. Nuclei were counterstained with DAPI (D9542, Sigma-Aldrich, 1:3000) and the slides were mounted with Fluoro-Gel (17985, Electron Microscopy Sciences). Primary antibodies used are described in Key Resources Table and secondary antibodies were Alexa Fluor-conjugated secondary antibodies (1:1000). EdU was detected using a Click-it EdU (C10340, Invitrogen) assay with Sulfo-Cyanine5 azide (A3330, Lumiprobe Corporation) following the manufacturer’s protocol.

### Image acquisition

Images were collected with a DM6000 Leica microscope, a NanoZoomer Digital Pathology microscope (Hamamatsu Photonics), or an LSM880 confocal microscope (Zeiss). Images were processed using NDP.view2 software, ImageJ software (NIH, Bethesda, MA, USA) and Photoshop (Adobe).

### Flow cytometry and Cell sorter

Cerebella were dissected into cold Hank’s Buffered Salt Solution (HBSS) (14170112, ThermoFisher) and then dissociated into Papain (LK003150, Worthington) at 37°C for 30 min. After dissociation, cells are centrifuged 5 minutes (min) at 400 g. Papain was inactivated with Ovomucoid protease inhibitor. After centrifugation, cells were finally washed in PBS filtered using a 40 μm mesh cell strainer (CLS431750, Sigma-Aldrich). After 5 min of centrifugation at 400g, the pellet was resuspended in PBS, 0.5% BSA for downstream experimentation.

### RT qPCR

The 4 populations of cells (GFP^-^ CFP^-^, GFP^+^ CFP^-^, GFP^-^ CFP^+^ and GFP^+^ CFP^+^) from *Nes-CFP*; *Atoh1-GFP* mice at P1 were sorted and frozen in TRI-reagent (Sigma-Aldrich). Total RNA was extracted using the mRNeasy kit (Qiagen), and cDNA was synthesized using SuperScript IV reverse transcriptase (Invitrogen). For the real-time PCR, 20ng of cDNA was used per reaction and amplified using StepOnePlus™ Real-Time PCR System. The reactions used a Step One Plus apparatus and software. The fold change was determined using the formula 2^-ΔΔCT^, where ΔΔCT= ΔCT_sample_ – ΔCT_GFP-CFP- sample_ with ΔCT= CT*_gene_* - CT*_Gapdh_*. Statistics were determined using the Mann-Whitney test. Sequence of primer pairs used are listed in the Key resources Table.

### Single-cell RNA sequence sample preparation and analysis

Single Cell RNA-seq sample preparation: *Nes-CFP* and *Atoh1-GFP*; *Nes-CFP* mice were euthanized at P1 and the cerebella were dissected in cold HBSS. For *Nes-CFP* cerebella, only the posterior part was kept, posterior to the primary fissure. Posterior cerebella or whole cerebella (*Atoh1-GFP*; *Nes-CFP* mice) were dissociated in Papain for 30 minutes at 37°C before being washed out in Ovalbumin Tumors were triturated by pipetting up and down using a P1000 and filtered through a 40µm cell strainer. Once the cells were dissociated, *Nes-CFP* cell pellets were washed with PBS. For each sample (4 samples, 1 cerebellum each), cells were counted using 0.2% Trypan on a hemocytometer and diluted before loading onto a 10X chip, targeting 10,000 viable cells per sample. *Atoh1-GFP*; *Nes-CFP* cell pellets were grouped together (3 cerebella for 1 sample) and the double positive population CFP^+^ GFP^+^ were sorted using BD FACS aria. Transgene negative, *Nes-CFP* and *Atoh1-GFP* cerebella were used as controls to set up the FACS settings. The scRNA-Seq of cell suspensions for the 5 samples was performed on a Chromium instrument (10X genomics) following the user guide manual for 3’ v3.1. Final libraries were sequenced on an Illumina NovaSeq6000 by the IGO core of MSKCC.

scRNA-seq data analysis: The Cell Ranger Single Cell software suite (10x Genomics) was used to align reads and generate feature-barcode matrices. The Genome Reference Consortium Mouse Build 38 (GRCm38, Gencode annotation mm10) was used as the reference genome. Raw reads were treated using the Cell Ranger count program with the default parameters. Seurat v5.0.2 package was used to generate a UMI (unique molecular identifier) count matrix from the Cell Ranger output38. Genes expressed in less than 10 cells were removed for further analyses. Cells with low transcriptomic complexity were filtered out by computing the ratio of detected genes to unique molecular identifiers (log10GenesPerUMI). Only cells with a log10GenesPerUMI > 0.80 were retained for downstream analyses to exclude low-quality or low-complexity cells. Furthermore, cells with less than 500 UMIs, more than 0.20 mitoRatio, and less than 250 genes detected were considered low quality/outliers and discarded from the datasets. Normalization was performed on individual samples using the NormalizeData function with default parameters. The normalized data was scaled by ScaleData function. Samples were then integrated using IntegrateData. The first 40 dimensions were used for the FindNeighbors function, and clusters were identified using the FindClusters function with a resolution of 0.8. Data were projected onto the 2D space using the FindUMAP or RunUMAP function with 40 dimensions. Cluster markers and further differential gene expression analyses were performed using normalized counts (NormalizeData) in the RNA assay. Cluster markers were identified using the FindAllMarkers and comparing markers generated to the literature. To refine clustering further, the SubsetData function was used to create a new Seurat object, and the above clustering was reiterated. Differential gene expression analyses were performed using the FindMarkers function in the Seurat package. Genes with an adjusted P value (Padj) < 0.05 were considered significantly up or downregulated. Results were visualized by Feature plot, Violin plot and Umap plot using Seurat package, and Volcano plot using the EnhancedVolcano package on Bioconductor39. GSEA analysis on GO and Hallmark terms were performed using the enrichGO, clusterProfiler packages on Bioconductor40.

### Quantification and statistical analysis

At least 3 mice of each genotype were used for all quantifications, and the numbers for each experiment are stated in the figure legends. A statistical method was not used to predetermine each sample size – the sample sizes are similar to previously reported work.

### Statistical analyses

All statistical analyses used to quantify data were performed using Prism software (GraphPad). Statistical significance was defined as p ≤ 0.05. The specific statistical tests applied are detailed in the corresponding figure legends. Data, presented as scatter plots, are shown as mean ± standard error of the mean (SEM).

## Acknowledgements

We thank members of the Joyner lab for interesting discussions, in particular Reeti Mayur Sanghrajka for insightful discussions and advice at the beginning of the project. We thank Christo Goridis for the *Atoh1^FRT-Cre^* mice and the Memorial Sloan Kettering Cancer Center Flow Cytometry Core and the Integrated Genomics Operations Core Facility for technical support.

## Additional Information

### Funding

NCI (R01CA192176), Alexandra L Joyner

NINDS (R01NS092096), Alexandra Joyner

Cycle for Survival, Alexandra L Joyner

NCI Cancer Center Support Grant (CCSG, P30 CA08748), Alexandra L Joyner

François Wallace Monahan Fellowship, Salsabiel El Nagar

### Ethics

There are no ethical considerations since no human data were used.

### Data Availability

Sequencing data are deposited in GEO (token inoxueqgzrqntgt) under accession code GSE328459. All other experimental data in this study are included in the main text, figures and graphs and Source data files in the manuscript. For the research, no new materials were generated.

## Supplementary Files

**Figure 2–Source data 1.** Source Data 1. RT-qPCR raw data and computation of ΔCT and 2-ΔΔCT values for comparing GFP^-^ CFP^+^, GFP^+^ CFP^-^ and GFP^+^ CFP^+^ populations to GFP^-^ CFP^-^population. Related to Figure 2C, D.

**Figure 5-Source data 1.** Gene expression in each cluster of the integrated 5 sample scRNA-seq data. Related to Figure 5A and Figure 5 supplement 1C.

**Figure 5-Source data 2.** Gene expression in each GCP cluster. Related to Figure 5D-I and Figure 5 supplement 1E

**Figure 5-Source data 3.** Cell cycle phase assignment for each cell of GCP clusters. Related to Figure 5J.

**Figure 5 - Source data 4.** ScRNA-seq significant differential gene expression analysis in cluster 10 compared to Clusters 1, 3 and 6 using FindMarkers approach.

**Figure 5-Source data 5.** ScRNA-seq significant differential gene expression analysis in cluster 10 compared to Clusters 1, 3 and 6. Related to Figure 5L

**Figure 5-Source data 6.** Gene Ontology (GO) analysis of genes upregulated in cluster 10 compared to clusters 1, 3 and 6. Related to Figure 5M

**Figure 5-Source data 7.** Gene Ontology (GO) analysis of genes downregulated in cluster 10 compared to clusters 1, 3 and 6. Related to Figure 5N

**Figure 6-Source data 1.** Gene expression in each cluster of integrated GCP clusters 1, 3, 6 and 10 and SmoM2 GCP-like tumor cells (Lao et al., 2026). Related to Figure 6A

**Figure 6-Source data 2.** ScRNA-seq significant differential gene expression analysis of GCP clusters 1, 3, 6 and 10 compared to SmoM2 GCP-like tumor cells. Related to Figure 6D

**Figure 6-Source data 3.** ScRNA-seq significant differential gene expression analysis of *Nes*-GCPs compared to SmoM2 GCP-like tumor cells. Related to Figure 6F

**Figure 6-Source data 4.** ScRNA-seq significant differential gene expression analysis of *Nes*^+^ GCPs compared to SmoM2 GCP-like tumor cells. Related to Figure 6H

---

**Figure 4 – figure supplement 1.**
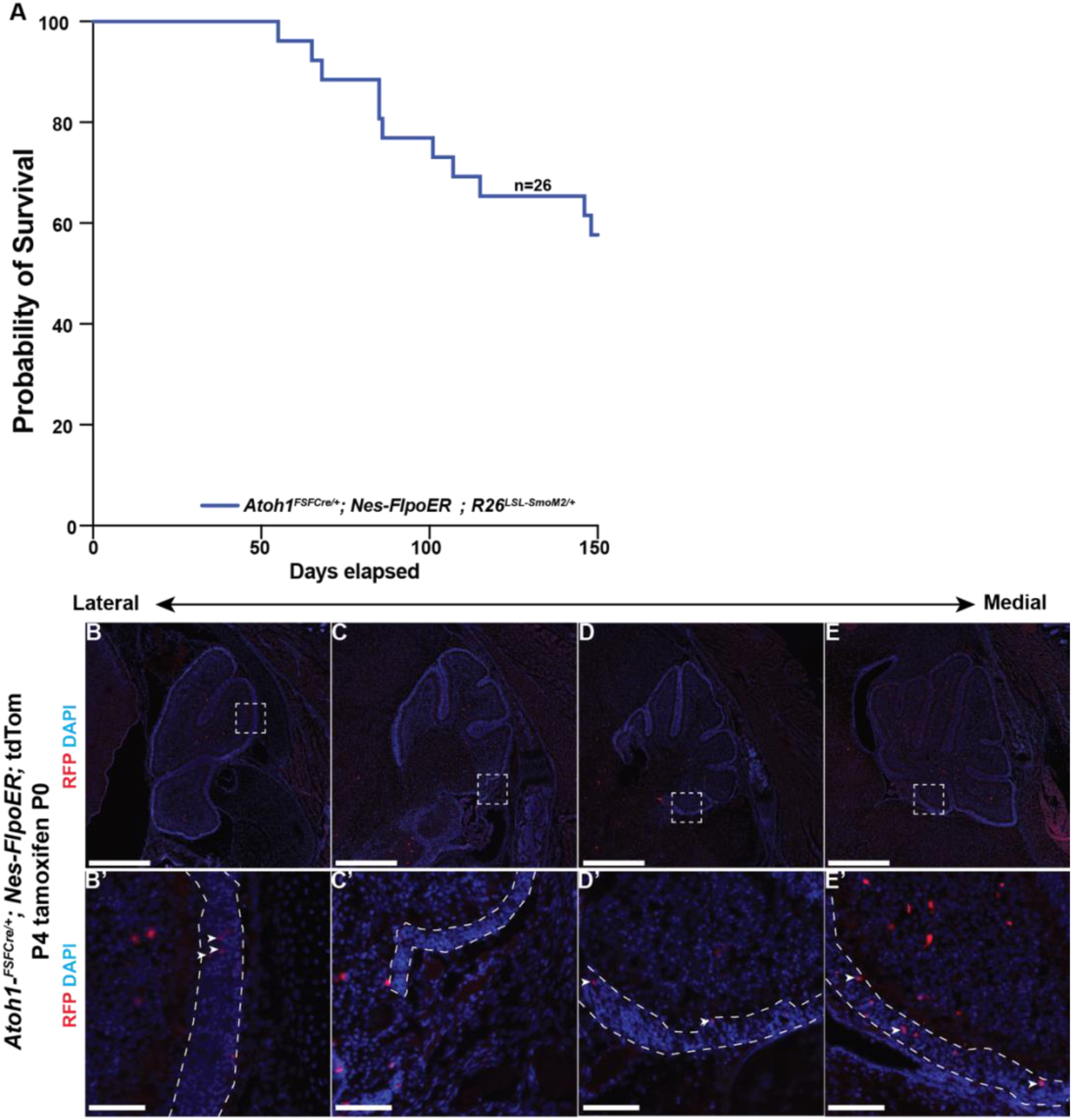
A dual recombinase strategy confirms SHH MB tumors can initiate from *Nes*-expressing GCPs (*Atoh1*^+^). **(A)** Kaplan-Meier survival curve of *Atoh1^Frt-Cre/+^; Nes-FlpoER; R26^LSL-SsmoM2/+^* mice following tamoxifen induction at P0. **(B-E)** Immunofluorescence staining of sagittal cerebellar sections from *Atoh1^Frt-Cre/+^; Nes-FlpoER;* tdTom mice collected at P4 following tamoxifen injection at P0 showing RFP (red) and DAPI (blue). Sections are shown at four distinct lateral-medial levels. (B’-E’) High magnification images of boxed regions in Panels (B-E). White arrowheads indicate the tdTomato^+^ cells (RFP) in the EGL. N>3 mouse samples. Scale bars: B-E: 500um, B’-E’: 50um

**Figure 4 – figure supplement 2.**
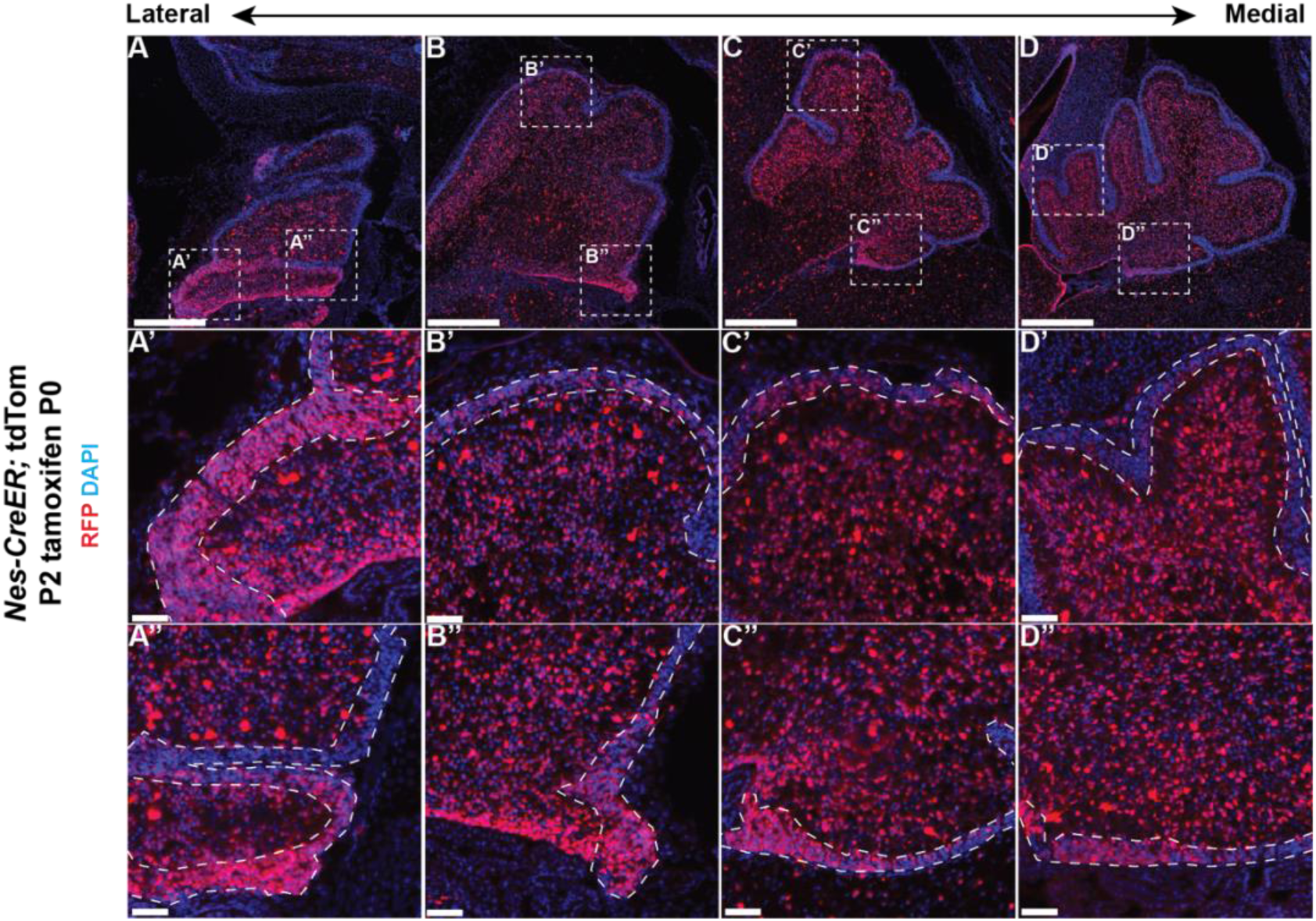
The *Nes-CreER* line labels a much broader progenitor population of cells than the *Nes-FlpoER* line. (A-D) Immunofluorescence staining of sagittal cerebellar sections from *Nes-CreER;* tdTom mice (*Nes-CreER; R26^LSL-tdTom^*) collected at P2 following tamoxifen injection at P0, showing RFP (red) and DAPI (blue). Sections are shown at four distinct lateral-medial levels. (A’-D’) High magnification images of anterior cerebellum. (A’’-D’’) High magnification images of posterior cerebellum. N= 3 mice. Scale bars: A-D: 500um and A’-D’, A’’-D’’: um

**Figure 5 – figure supplement 1.**
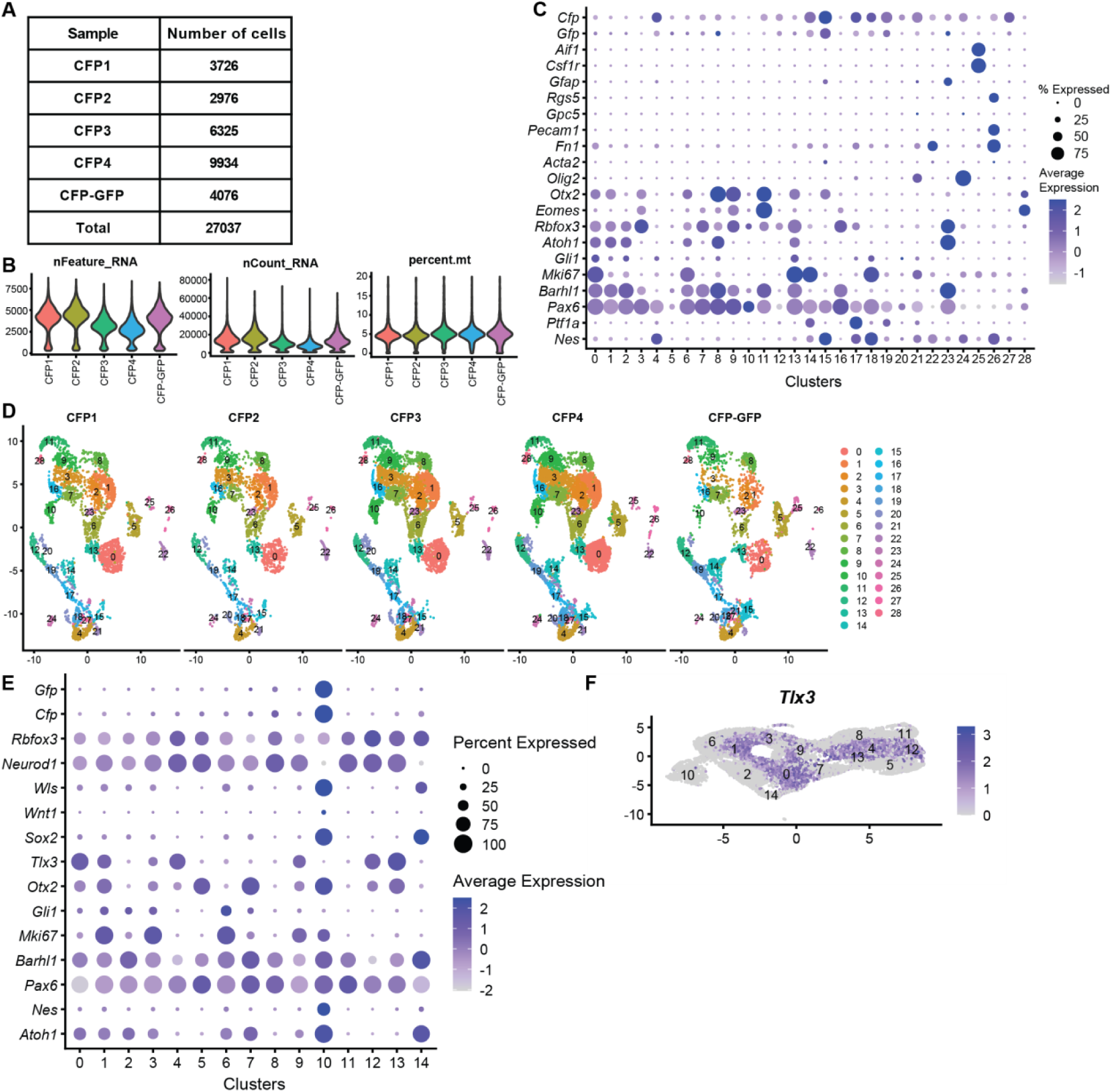
ScRNA-seq workflow and additional characterization of the *Nes*-expressing GCP clusters. **(A)** Number of cells from each sample used for downstream analyses after filtering. Samples CFP1 to CFP4 represent 1 animal each and CFP-GFP represent 3 animals pooled. **(B)** Violin plots showing the number of features, the RNA counts and the percent of mitochondrial RNA counts in each sample. **(C)** Dot plot graph showing expression levels of cell type marker genes across all clusters for all cells. See also Figure 5-Source data 1. **(D)** UMAPs of all cells shown by samples. CFP1 to CFP4 are the posterior GCP samples from *Nes-CFP* mice (one mouse per sample) at P1 and CFP-GFP corresponds to the sample from *Atoh1-GFP*; *Nes-CFP* mice where CFP^+^ GFP^+^ cells were isolated by FACS (n=3 mice combined). **(E)** Dot plot graph showing expression levels of GCP markers across the integrated GCP dataset. See also Figure 5-Source data 2. **(F)** Feature plots showing expression of posterior marker *Tlx3* in GCP clusters in the integrated GCP dataset. See also Figure 5-Source data 2.

